# The Effects of Vitamin E Analogues α-Tocopherol and γ-Tocotrienol on the Human Osteocyte Response to Ultra-high Molecular Weight Polyethylene Wear Particles

**DOI:** 10.1101/2022.04.17.488608

**Authors:** Renee T Ormsby, Kunihiro Hosaka, Andreas Evdokiou, Andreani Odysseos, David M Findlay, Lucian B. Solomon, Gerald J Atkins

## Abstract

Polyethylene (PE) liners are a common bearing surface of orthopaedic prostheses. Wear particles of ultra-high molecular weight PE (UHMWPE) contribute to periprosthetic osteolysis, a major cause of aseptic loosening. Vitamin E is added to some PE liners to prevent oxidative degradation. Osteocytes, an important cell type for controlling both bone mineralisation and bone resorption, have been shown to respond UHMWPE particles by upregulating pro-osteoclastogenic and osteocytic osteolysis. Here, we examined the effects of the vitamin E analogues α-tocopherol and γ-tocotrienol alone or in the context of UHMWPE particles on human osteocyte gene expression and mineralisation behaviour. Human osteoblasts differentiated to an osteocyte-like stage were exposed to UHMWPE wear particles in the presence or absence of either α-Tocopherol or γ-Tocotrienol. Both α-Tocopherol and γ-Tocotrienol induced antioxidant-related gene expression. UHMWPE particles independently upregulated antioxidant gene expression, suggesting an effect of wear particles on oxidative stress. Both vitamin E analogues strongly induced *OPG* mRNA expression and γ-Tocotrienol also inhibited *RANKL* mRNA expression, resulting in a significantly reduced *RANKL*:*OPG* mRNA ratio (p < 0.01) overall. UHMWPE particles reversed the suppressive effect of α-Tocopherol but not of γ-Tocotrienol on this pro-osteoclastogenic index. UHMWPE particles also upregulated osteocytic-osteolysis related gene expression. Vitamin E analogues alone or in combination with UHMWPE particles also resulted in upregulation of these genes. Consistent with this, both vitamin E analogues promoted calcium release from mineralised cultures of osteocyte-like cells. Our findings suggest that while vitamin E may suppress osteocyte support of osteoclastogenesis in the presence of UHMWPE particles, the antioxidant effect may induce osteocytic osteolysis, which could promote periprosthetic osteolysis. It will be important to conduct further studies of vitamin E to determine the long-term effects of its inclusion in prosthetic materials.

## Introduction

Implant loosening is a major problem associated with articulating joint replacement surgery. A major cause of aseptic loosening is the generation of wear particles of implant materials leading to the development of peri-prosthetic osteolytic lesions. Ultra-high molecular weight polyethylene (UHMWPE) is a common liner used for the bearing surface and wear particles of this are directly associated with the production of osteolytic lesions, with high volumetric wear rates correlating with the greatest progression of lesion volume (1). UHMWPE liner wear due to mechanical friction against the metal femoral head, may be exacerbated by oxidative degradation of the polyethylene (PE). UHMWPE wear particles are known to stimulate a variety of cellular pathways, including osteoclastic bone resorption, inflammation, inhibition of bone formation (2), as well as osteocytic peri-lacunocanalicular remodelling, also known as osteocytic osteolysis (3).

The addition of Vitamin E analogues is a recent approach to improving the endurance of UHMWPE liners by preventing oxidation. The aim of imbuing PE liners with Vitamin E is to reduce the formation of free radicals within the PE, as well as improving fatigue properties that would normally occur after post-irradiation melting, without sacrificing fatigue strength (4). Vitamin E, specifically α-tocopherol, can be blended with UHMWPE powder prior to radiation, which protects against oxidation within the PE chain but reduces the crosslinking efficiency (5–7). Alternatively, UHMWPE is diffused with vitamin E after radiation, circumventing possible changes to the crosslinking efficiency (8). However, diffusion of vitamin E would result in relatively less vitamin E covalently bound within the PE chain compared to addition prior to radiation, potentially allowing unbound vitamin E to leach into the synovial fluid and surrounding tissues, including bone tissue (9).

Both Tocopherols and Tocotrienols play potentially important roles in regulating bone resorption (10, 11). α-Tocopherol has been shown to have varying effects on both bone formation and resorption, dependent on the dose and model system investigated (10). For example, specific knockdown of the α-tocopherol transfer protein in mice, where vitamin E absorption is blocked, showed a significantly higher bone mass due to lower bone resorption and reduced osteoclast surface (12). Tocotrienol is able to reduce oxidative damage in osteoblasts, promoting osteoblast differentiation and survival (13). Human epidemiological studies have also investigated the relationship between vitamin E and bone metabolism, however, with mixed results. One study, in a US population aged 50 and older, described a negative correlation between the high serum concentration of tocopherol and the femoral neck BMD (14). Similarly, a study by Wolf *et al*. showed a negative correlation between total vitamin E intake and femoral neck BMD in an age-adjusted regression analysis of women in the USA (15). However, a longitudinal study on Swedish subjects over the age of 50 and monitored for 19 years, has shown an association between low intake and serum concentration of α-tocopherol, and an increased rate of fracture (16).

Osteocytes are the most abundant cell type in bone and are critical in controlling bone remodelling and maintaining skeletal integrity (17). Osteocytes are capable of resorbing their perilacunar matrix by producing bone degrading enzymes, including matrix metallopeptidase 13 (*MMP13*) (18, 19), Cathepsin K (*CTSK*) (20) and carbonic anhydrase II (*CA2*) (21). Human osteocytes exposed to UHMWPE particles *in vitro* display both osteocytic osteolytic and pro-osteoclastogenic responses (3, 22). Examination of human bone biopsies from implant patients with aseptic loosening and with confirmed periprosthetic osteolysis, also showed significantly increased osteocyte lacunar area compared to those in primary THR biopsies (3, 22). Little is known regarding the effects of UHMWPE particles on bone cells, such as osteocytes, in the added presence of vitamin E. In this study, we investigated the biological effects of two major analogues of vitamin E, α-Tocopherol and γ-Tocotrienol, in the presence or absence of UHMWPE particles, on human primary osteocyte-like cells in terms of oxidative stress and bone catabolic responses.

## Materials and Methods

### Patient bone-derived cells

All patient-derived samples were obtained with written, informed consent and human research ethics approval (Royal Adelaide Hospital Human Research Ethics Committee approval no. 090101). Human primary osteoblasts (Normal Human Bone-derived Cells; NHBC) were derived from proximal femoral cancellous bone biopsies, taken from patients undergoing total hip replacement surgery, and cultured and passaged *ex vivo*, as described previously (19, 22).

### Osteocyte-like cell culture

NHBC isolated from 3 different donors were cultured for 28 days until the cells reached a mature osteocyte-like stage, as described previously (19, 22, 23). The cells were then treated for a further 7d with either α-Tocopherol or γ-Tocotrienol (Sigma Aldrich, St Louis, MO, USA) at concentrations of 100 nM, 300 nM, 1 μM or untreated control with experimental quadruplicates. Similarly, osteocyte-like cells after 28d of culture were overlaid with a collagen gel (Cellmatrix Type 1-A, Nitta, Tokyo, Japan) (3), containing UHMWPE particles (Ceridust 3615, Clariant Company) (100μg/ml), with the addition of either α-Tocopherol or γ-Tocotrienol (Sigma Aldrich, St Louis, MO, USA) (1 μM), or particle free and vehicle controls, and cultured for a further 21d. UHMWPE particles were confirmed to be endotoxin free and were sized from 0.3-10 μM. Either α-Tocopherol or γ-Tocotrienol, 1 μM was added to the culture medium every 3 days.

### Gene Expression

Total RNA was extracted using TRIzol (Life Technologies, NY, USA) and complementary DNA was synthesised using iScript™ RT kit (Bio-Rad, Hercules, CA, USA), according to the manufacturer’s instructions. Real time RT-PCR was performed using SYBR Green Fluor qPCR Mastermix (Qiagen, Limburg, The Netherlands) on a CFX Connect (Bio-Rad, Hercules, CA, USA). Gene expression was normalised to that of the housekeeping gene, *HPRT1*. Oligonucleotide sequences for the mRNA-specific amplification of *MMP13, CA2, CTSK, RANKL, OPG* and *CSF1* are published (3). Oligonucleotide sequences for *HPRT1, SOD1, SOD2* and *CAT* are listed in **Table 1**. All primers were designed in-house to be mRNA specific by at least one per pair flanking an intron-exon boundary, and all were purchased from Sigma Aldrich (St. Louis, MO, USA).

**Table 1:**
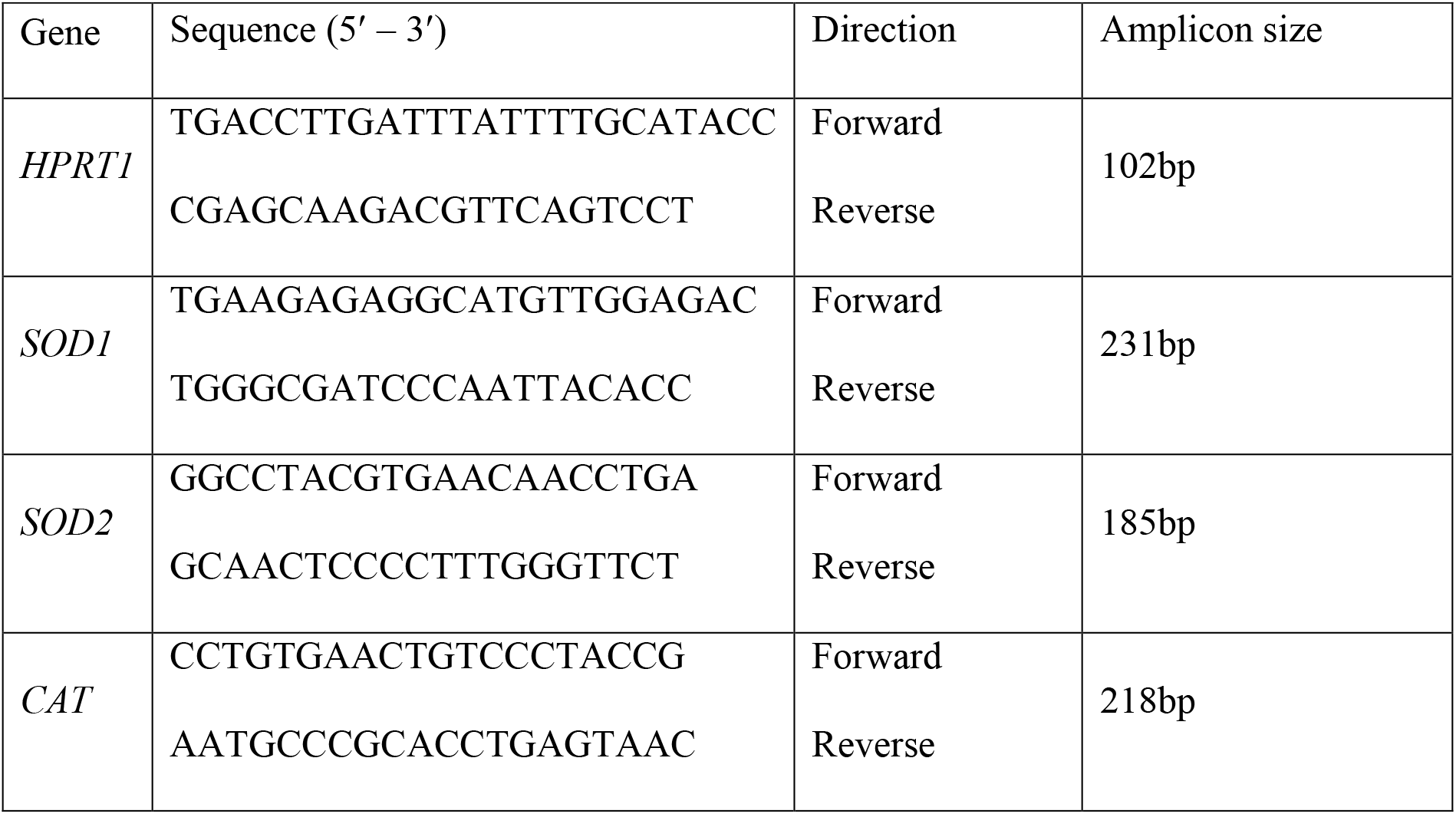
Sequences of sense (S) and anti-sense (AS)-specific oligonucleotide primers and predicted PCR product sizes. Primers were designed to be mRNA-specific.

### In vitro mineralisation

Calcium deposition was visualised using the Alizarin Red stain, as described previously (24). Briefly, quadruplicate wells per treatment were washed with PBS and then fixed with 10% neutral buffered formalin for 1 hour at RT and then washed with PBS (3x). Each well was stained with aqueous 2% (w/v) Alizarin red solution (Sigma Aldrich) at pH 4.2, for 5 minutes at room temperature and then washed with distilled water to remove any unbound stain. Each well was imaged (Olympus SZ2-ILST, Tokyo, Japan) and then the bound stain was solubilised with 100 μl of 10% Cetylpyrridium chloride (CPC) (Sigma Aldrich) in 10mM Sodium phosphate monobasic monohydrate (pH 7.0) added to each well for 15 minutes at RT. The optical density was determined at 560nm by spectrophotometry (Thermo Multiskan Ascent).

### Statistical analysis

Statistical differences within data sets were analysed using GraphPad Prism Version 9.0.0 software (GraphPad Prism, La Jolla, LA, USA). Differences in normalised gene expression to control values were analysed using non-parametric Kruskal-Wallis tests with Mann Whitney post-hoc tests. Differences in Alizarin Red mineralisation were tested using one-way analysis of variance (ANOVA) with Tukey’s multiple comparisons tests. In all cases a value for *p* < 0.05 was considered significant.

## Results and Discussion

### Effects of UHMWPE and Vitamin E analogues on the oxidative stress response

In order to investigate the effects of vitamin E analogues and UHMWPE particles, we used a clinically relevant model where human primary osteoblasts isolated from THR implant recipients, the population at risk from developing periprosthetic osteolysis, were cultured for 28 days under pro-osteogenic conditions to generate osteocyte-like cultures (3). Once established, the cells were overlaid with a type I collagen gel containing UHMWPE particles to promote cell-particle contact, with the addition of either α-Tocopherol or γ-Tocotrienol and cultured for a further 21 days.

Vitamin E is an attractive bone supplement due to its anti-inflammatory and antioxidant activities. The induction of antioxidant gene expression can be both a marker of oxidative stress and an indication of a physiological response to the causative reactive oxygen species (ROS). Therefore, we investigated the expression of key antioxidant genes, superoxide dismutases 1 and 2 (*SOD1* and *SOD2*), which are produced in response to ROS and act by binding to superoxide radicals, metabolising them to oxygen and hydrogen peroxide (25). The expression of catalase (*CAT*), which further converts hydrogen peroxide into water and oxygen (25), was also examined. Osteocyte-like cultures treated with either α-Tocopherol or γ-Tocotrienol demonstrated significantly upregulated expression of all three antioxidant genes (Fig. 1A-C). Moderate upregulation of these genes also occurred with exposure to UHMWPE particles alone, suggesting induction of oxidative stress. Importantly, both Vitamin E analogues stimulated further increased expression of these genes in the presence of UHMWPE particles. Previous studies have shown that PE liners and the resulting particles undergo oxidative degradation and can generate ROS (26, 27). Therefore, upregulation of antioxidant genes could be a protective mechanism by the osteocyte in response to ROS produced by the UHMWPE particles, preventing DNA damage and potential cell death. Consistent with this protective mechanism, we have previously reported that UHMWPE particles do not induce apoptosis in human osteocyte-like cells (3).

**Figure 1:**
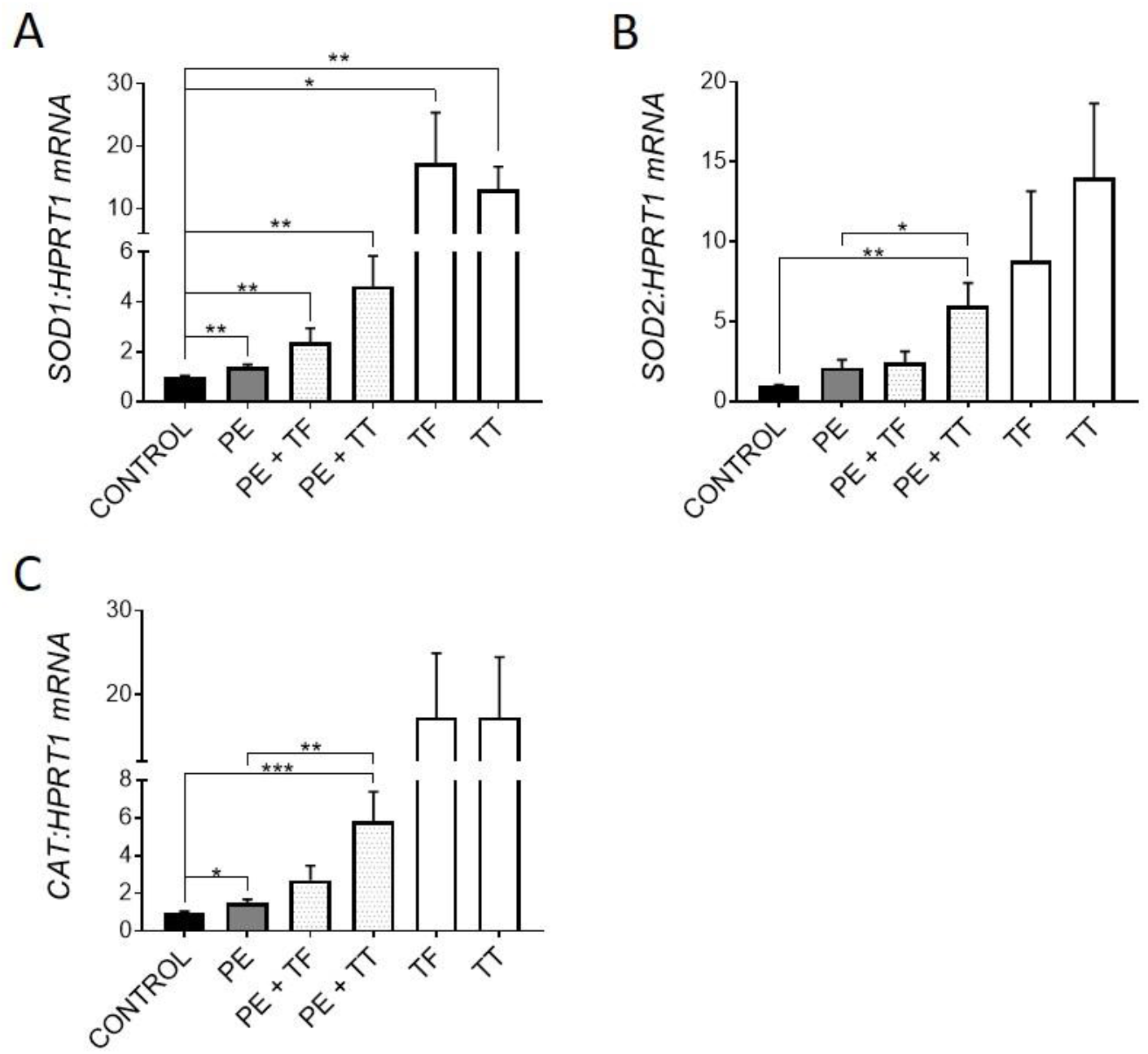
Effect of vitamin E and UHMWPE particles on antioxidant gene expression. Human osteocyte-like cultures were overlaid with collagen gel containing UHMWPE particles and either α-Tocopherol (TF) or γ-Tocotrienol (TT), and cultured for a further 21d. Relative gene expression was measured for antioxidant genes *SOD1*, *SOD2* and Catalase (A-C). Data shown are means ± standard error of the mean (SEM) of quadruplicate experiments analysed in duplicate. Significant differences are denoted by \**p* < 0.05, \*\**p* < 0.01, \*\*\**p* < 0.001.

### Effects of UHMWPE and Vitamin E on the pro-osteoclastogenic response

We next investigated the expression of genes associated with osteoclastogenesis. It was previously shown that osteocytes respond to UHMWPE wear particles by upregulating the pro-osteoclastogenic *RANKL:OPG* mRNA ratio, as well as upregulating key osteoclastic regulatory genes, *CSF1* and *IL8* (3, 28). Here, osteocytes responded to treatment with either α-Tocopherol and γ-Tocotrienol by downregulating the *RANKL:OPG* mRNA ratio. In particular, treatment with γ-Tocotrienol suppressed *RANKL* and increased *OPG* mRNA expression in the primary human osteocyte cultures (Fig. 2A-B), whereas α-Tocopherol, whilst having non-significant effects on either gene alone, also resulted in reduced *RANKL:OPG* expression overall.

**Figure 2:**
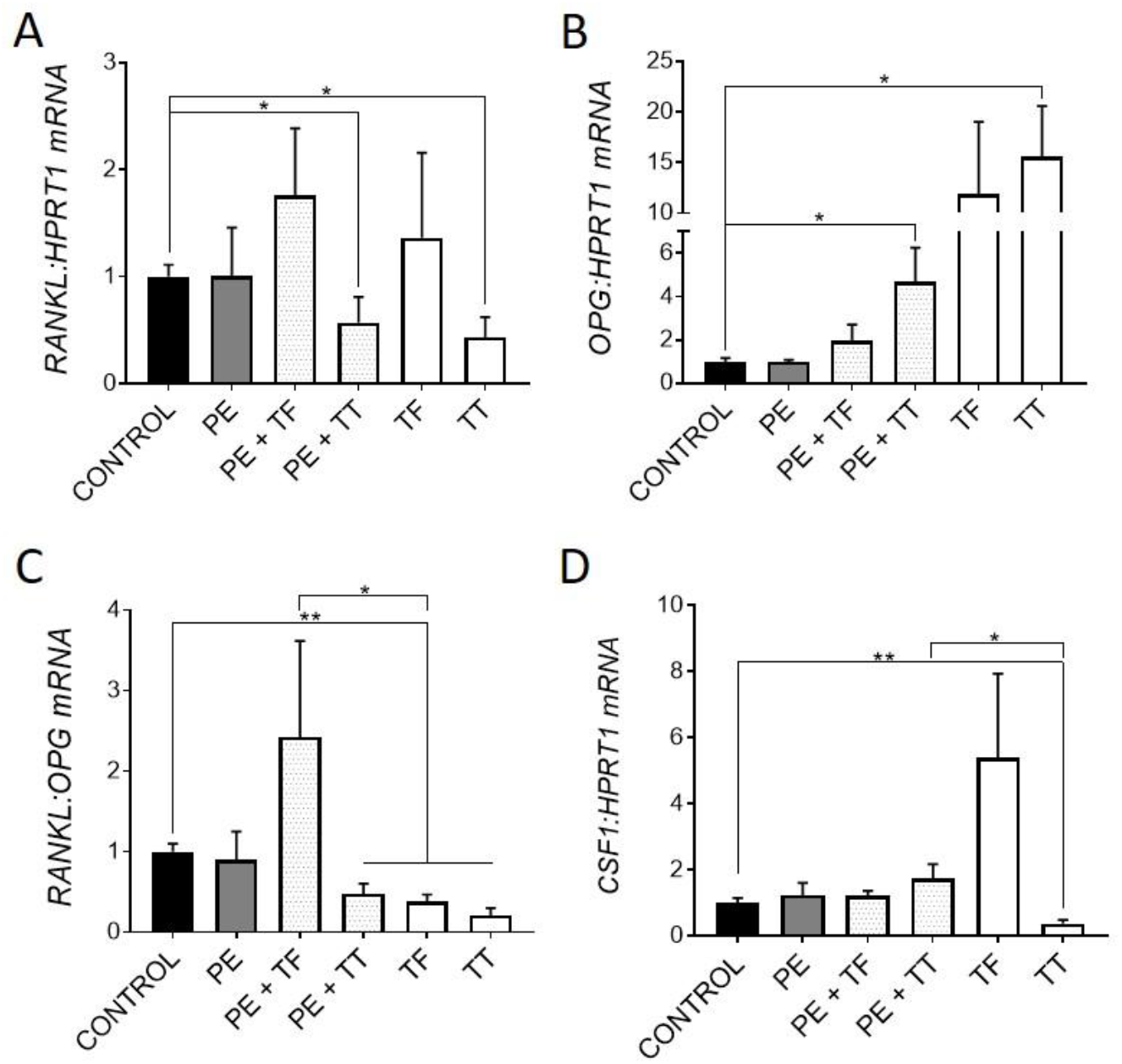
Effect of vitamin E and UHMWPE particles on osteoclastogenic gene expression. Relative gene expression was measured for genes associated with osteoclastogenesis: *RANKL*, *OPG*, the pro-osteoclastogenic *RANKL:OPG* mRNA ratio and *CSF1* mRNA expression (A-D). Data shown are means ± standard error of the mean (SEM) of quadruplicate experiments analysed in duplicate. Significant differences are denoted by \**p* < 0.05, \*\**p* < 0.01, \*\*\**p* < 0.001.

The addition of α-Tocopherol had no effect on the UHMWPE response in terms of this ratio. However, γ-Tocotrienol significantly reduced the *RANKL:OPG* mRNA ratio in the presence of particles. Interestingly, γ-Tocotrienol but not α-Tocopherol alone significantly downregulated *CSF1* expression, the protein product of which, CSF1/macrophage colony stimulating factor (M-CSF), is a key co-factor for RANKL-mediated osteoclastogenesis, although this effect was not seen in the presence of UHMWPE (Fig. 2D). Overall, these findings suggest that both α-Tocopherol and γ-Tocotrienol exert an anti-osteoclastogenic effect on human osteocyte-like cells, in the presence or absence of UHMWPE particles. Consistent with this, previous studies have shown that mice treated with vitamin E, either α-Tocopherol or γ-Tocotrienol, have decreased bone resorption (11, 29). Furthermore, a study by Bichara *et al*. (30) showed Vitamin E-bound UHMWPE wear particles significantly decreased the amount of bone resorption in a mouse calvarial model of osteolysis when compared to conventional UHMWPE particles.

### Effects of UHMWPE and Vitamin E on the osteocytic osteolysis response

Consistent with previous reports (3, 22), UHMWPE particles induced expression of genes associated with osteocytic osteolysis, including *MMP13, CA2* (not significantly) and *CTSK (p* < 0.001) (Fig. 3A-C).

**Figure 3:**
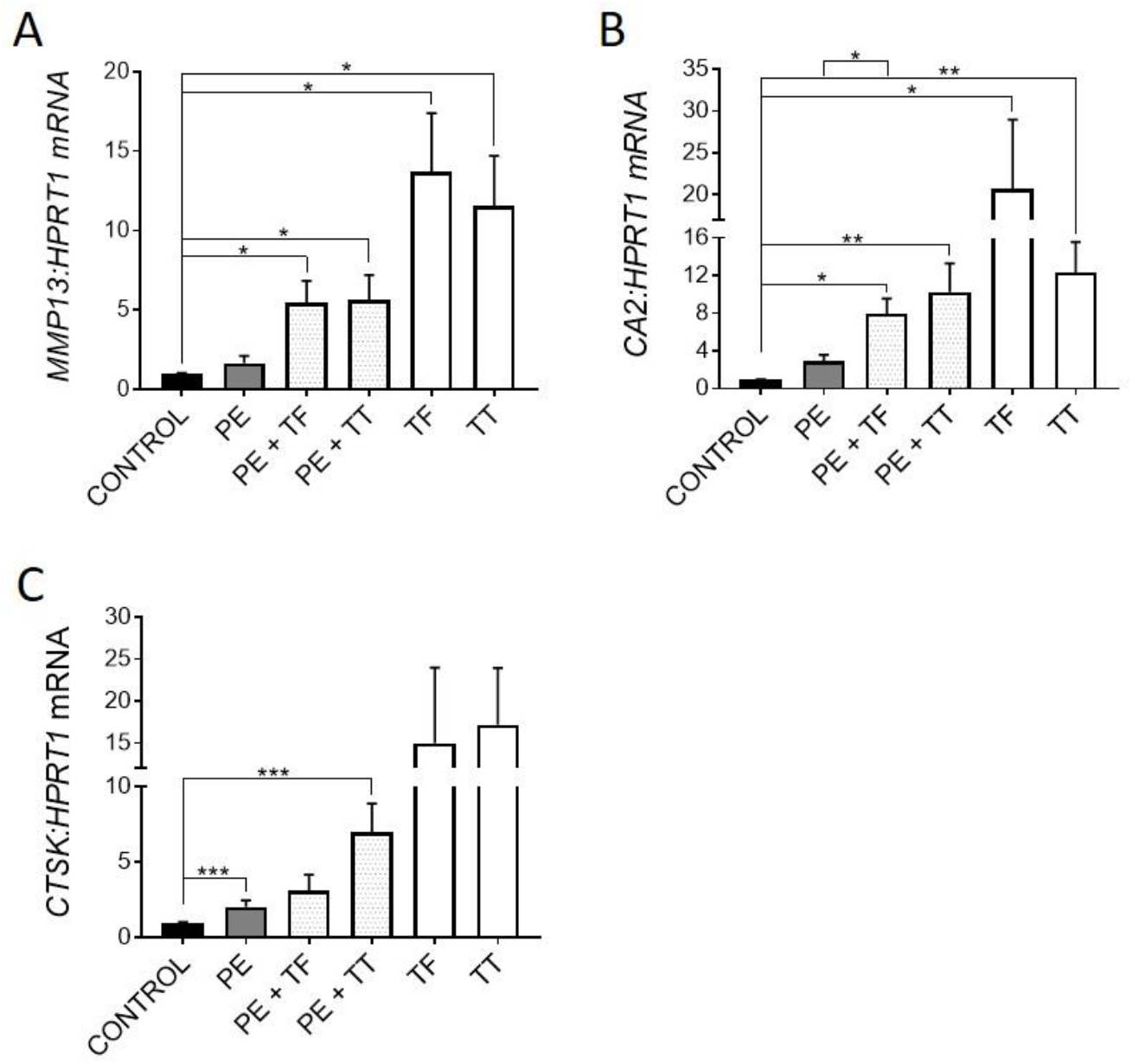
Effect of vitamin E and UHMWPE particles on expression of genes associated with osteocytic osteolysis. A. *MMP13* mRNA expression, B. *CA2* mRNA expression, C. *CTSK* mRNA expression. Data shown are means ± standard error of the mean (SEM) of quadruplicate experiments analysed in duplicate. Significant differences are denoted by \**p* < 0.05, \*\**p* < 0.01, \*\*\**p* < 0.001.

Treatment with either Vitamin E analogue upregulated the expression of all three genes, suggesting that Vitamin E could promote osteocytic osteolysis. Both α-Tocopherol and γ-Tocotrienol were able to further increase the expression of these genes in the presence of PE particles. As mentioned previously, UHMWPE particles are capable of producing ROS, which play a key role in osteoclastic differentiation and activity (31). Therefore, here, osteocytes could be responding to ROS through one of two pathways. Firstly, the inhibition of ROS in human osteocytes by treatment with Vitamin E analogues alone and in the presence of UHMWPE particles may have caused the increase in expression of genes associated with osteocytic osteolysis, *MMP13, CTSK* and *CA2*. Secondly, UHMWPE particles may stimulate the production of ROS by the osteocyte, inducing oxidative stress, triggering a perilacunar remodelling response. This effect would explain why there was upregulation of antioxidant gene expression in response to UHMWPE. Therefore, removal of perilacunar bone could be one of the mechanisms used by osteocytes to relieve oxidative stress. Further investigation is required to determine if ROS produced by the osteocyte plays a role in regulating osteocytic osteolysis *in vivo*.

### Effects of Vitamin E analogues on mineral handling

To investigate the effects of α-Tocopherol and γ-Tocotrienol on osteocytic regulation of mineralisation, osteocyte-like cultures were exposed to either analogue (0, 100 nM, 300 nM and 1 μM) in the absence of UHMWPE particles. Calcium deposition was measured using the Alizarin Red stain, as described previously (24). Consistent with the observed regulation of osteocytic osteolysis related genes, treatment with either Vitamin E analogue resulted in decreased mineralisation (Fig. 4).

**Figure 4:**
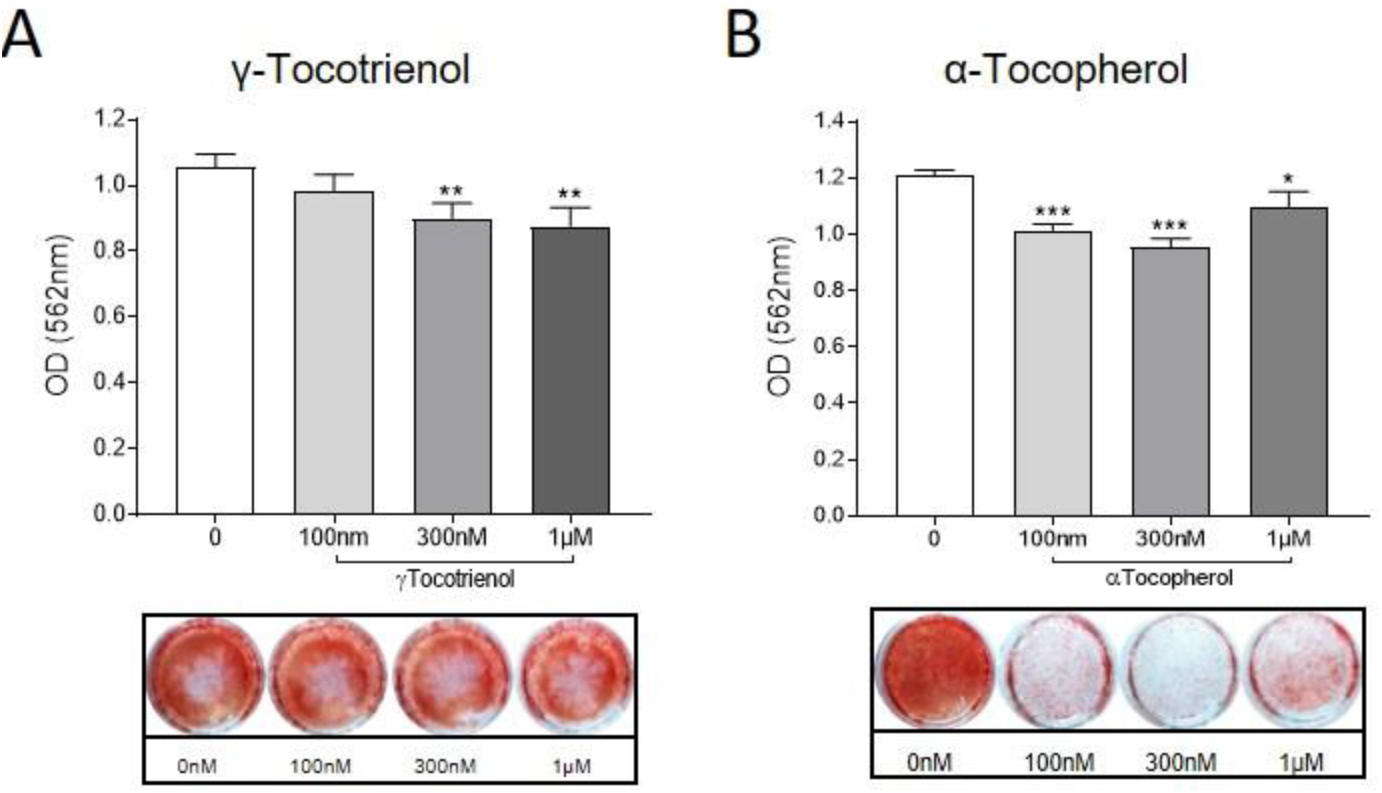
Effect of vitamin E analogues on osteocyte mineral handling. Human osteocyte-like cultures were treated with either α-Tocopherol (TF) or γ-Tocotrienol (100nM, 300nM, 1μM) or untreated (0) for a further 6d. Cells were stained with Alizarin Red and then quantified using Cetylpyrridium chloride to elute the bound stain. Absorbance at 560nM was measured by spectrophotometry. Data shown are means ± standard deviation of quadruplicate wells. Significant differences are denoted by \**p* < 0.05, \*\**p* < 0.01, \*\*\**p* < 0.001.

This suggests that vitamin E could contribute to the loss of bone that occurs during periprosthetic osteolysis specifically through promoting osteocytic removal of perilacunar bone (21).

## Conclusions

This study demonstrates the potential effects of α-Tocopherol and γ-Tocotrienol on the regulation of bone resorption by the human osteocyte. Our data suggest that UHMWPE particles induce oxidative stress in osteocytes, which could be a potential trigger of perilacunar remodelling. Two distinct actions of the vitamin E analogues were observed: suppression of the pro-osteoclastic response, and stimulation of the osteocytic osteolysis response. Thus, while our findings are consistent with vitamin E suppressing the osteoclastic response to UHMWPE particles, it could exacerbate an osteocytic osteolysis response, which in turn could contribute to periprosthetic osteolysis. The effects of vitamin E analogues on osteocytes requires further investigation to determine if their lytic effects have potential consequences in the development of periprosthetic osteolysis and aseptic loosening.

## Acknowledgements

The authors thank the nursing and surgical staff of the Orthopaedic Trauma Service, RAH, for their help in collecting patient bone specimens. This work was supported by funding from the National Health and Medical Research Council of Australia (NHMRC) Project Grant Scheme (Grant ID 1041456) and the Royal Adelaide Hospital Special Purposes Fund. RTO was the recipient of a University of Adelaide post-graduate scholarship. GJA was supported by a NHMRC Senior Research Fellow fellowship.

